# Widespread Suppression of High-Order Visual Cortex During Blinks and External Predictable Visual Interruptions

**DOI:** 10.1101/456566

**Authors:** Tal Golan, Shany Grossman, Leon Y Deouell, Rafael Malach

## Abstract

Spontaneous eye blinks generate frequent potent interruptions to the retinal input and yet go unnoticed. As such, they provide an attractive approach to the study of the neural correlates of visual awareness. Here, we tested the potential role of predictability in generating blink-related effects using fMRI. While participants attentively watched still images of faces and houses, we monitored naturally occurring spontaneous blinks and introduced three kinds of matched visual interruptions: cued voluntary blinks, self-initiated (and hence, predictable) external darkenings, and physically similar but unpredictable external darkenings. These events’ impact was inspected using fMRI across the visual hierarchy. In early visual cortex, both spontaneous and voluntary blinks, as well as predictable and unpredictable external darkenings, led to largely similar positive responses in peripheral representations. In mid- and high-level visual cortex, all predictable conditions (spontaneous blinks, voluntary blinks, and self-initiated external darkenings) were associated with signal decreases. In contrast, unpredictable darkenings were associated with signal increases. These findings suggest that general-purpose prediction-related mechanisms are involved in producing a small but widespread suppression of mid- and high-order visual regions during blinks. Such suppression may down-regulate responses to predictable transients in the human visual hierarchy.

---

When taking a photograph of a large group of people, almost inevitably at least one of them will be photographed with their eyes closed due to spontaneous eye blinking. In a letter to Nature, Lawson (1948) has counted such unfortunate closed eyes photographees, combining this proportion with estimates of spontaneous blink rates to obtain estimates of the duration during which an eye blink shuts vision – 270-330 ms. Later estimates of this ‘blackout duration’ are shorter (100-150 ms, see Riggs et al. 1981) but it is still striking that these frequent eye closures somehow evade our awareness; external darkenings of a matched duration are easily seen (Riggs et al. 1981).

The question of how the human visual system deals with the frequent interruption caused by eye blinks has attracted considerable scientific attention. Volkmann and colleagues (1980) showed that during voluntary blinks, psychophysical sensitivity to flickering light emitted through the palate (hence bypassing the eyelids) is mildly suppressed, an effect which can be explained only by an extra-retinal mechanism. 25 years later, reapplying flickering light through the palate inside an MRI scanner, Bristow and Rees (2005) have shown that voluntary blinking (which unlike spontaneous blinks does produce a noticeable darkening) is associated with suppression of early visual fMRI BOLD response, an effect increasing in strength from V1 to V3.

An additional human fMRI study on the effect of blinks on visual cortex responses found that voluntary blinks reduced the response to contrast-reversing checkerboards more than matched external darkenings, an effect observed in several extra-striate visual regions, including MT (Bristow, Frith, et al. 2005). fMRI studies on spontaneous blinks found blink-related activation of the peripheral fields of V1 to V3 (Tse et al. 2010; Hupé et al. 2012). Spontaneous blinks during movie watching were found to be correlated with default mode activation (Nakano et al. 2013). Finally, our recent intracranial EEG (iEEG) study of spontaneous blinks (Golan et al. 2016) revealed that positive stimulus-driven activation transients in high order visual areas were suppressed during spontaneous blinks, potentially accounting for the lack of visual awareness of these interruptions.

A central question that remains unanswered concerns the mechanism underlying the suppression of eye blink-related neural responses. It is often assumed that suppression of blink-related activity is a product of a ‘hardwired’ efferent copy relayed from the oculomotor system to the visual cortex (e.g., Riggs *et al.* 1981). However, it is possible that more-general, flexible, mechanisms are involved as well. In particular, when eye blinks and external darkenings are contrasted, at least one major general cognitive factor confounds this contrast – predictability. Whereas eye blinks (either spontaneous or voluntary) are self-initiated and thus predictable, external darkenings, as applied in all of the studies that used this control (including Gawne and Martin 2000, 2002; Bristow, Frith, *et al.* 2005; Nakano *et al.* 2013; Golan *et al.* 2016), were controlled by the experimenter and presented in an unpredictable manner.

Thus, it might be that the differential physiological and behavioral responses to blinks and external darkenings reported in the above studies were not caused only by a specialized, hardwired efferent copy mechanism but may also involve a non-specific predictive-coding based error-signal, representing the surprising aspect of the external darkenings. This hypothesis predicts that darkenings rendered predictable by coupling their presentation to a self-generated motor-action (i.e., a button press) would reproduce at least some of the blink-related responses. To examine this possibility, we used fMRI to test the hemodynamic responses in the visual cortex to four brief events: spontaneous eye blinks, voluntary eye blinks, self-initiated darkenings - mimicking the motor-sensory coupling of blinks through forming an arbitrary motor-sensory link, and externally induced darkenings - mimicking the retinal impact of blinks but presented in an unpredictable manner. Each of the self-initiated darkenings was triggered by a button press executed by the subject in response to an auditory cue. All of the four different visual events were introduced in a matched fashion while the participants were attentively observing photographs of faces or houses.

In brief, we found that an extensive portion of mid- to high-level visual regions showed a small but significant reduction in their activity in response to all predictable interruptions: spontaneous blinks, voluntary blinks, and self-initiated darkening. In contrast, unpredictable external darkenings triggered a positive response in virtually all visual regions. The results indicate that in addition to the specialized hardwired blink-suppression pathway, general-purpose, more flexible predictive-computation mechanisms may participate in generating the blink-related neural suppression of high-level visual cortex.

## Materials and Methods

### Participants

13 healthy volunteers (8 females; mean age = 27.7, SD = 3.5; 12 right handed) participated in the fMRI experiment. The number of subjects was predetermined and was not conditioned on the results. Twelve of the participants underwent two scanning sessions, and one participant underwent one scanning session. All participants had normal vision, provided prior written consent, and were paid for their participation in the study. The Tel Aviv Sourasky Medical Center Institutional Review Board (Helsinki Committee) approved the procedures.

### Preliminary training

Since voluntary blinks tend to have a greater duration than spontaneous blinks (e.g., Sun et al. 1997), we trained the participants to adjust their voluntary blinks so their duration will match the average duration of their spontaneous blinks. This training was held in a room in the vicinity of the MRI scanner before each scanning session. First, we presented the participants with six experimental blocks similar to those used in the main experiment. The durations of the spontaneous blinks occurring during these blocks were recorded by a video eye-tracker (EyeLink 1000, SR research, Ontario, Canada). Blink detection and duration measurement were based on the eye tracker’s native eye blink detection algorithm (but see ‘blink detection and duration measurement’ below for improved detection and duration estimation used in the main experiment).

Next, a blank screen with a fixation mark was presented, and the participants had to blink once each time an auditory cue was given. Each such voluntary blink was followed by visual feedback consisting of a horizontal line with two vertical bars marking the desired range of voluntary blinks’ duration (determined to be ±1 SD of the participant’s mean spontaneous blinks duration), and a dot marking the duration of the participant’s most recent voluntary blink. Training was completed after five successive successful trials, i.e., five successive voluntary blinks that fell within the range of one standard deviation from the mean duration of spontaneous blinks. This required no more than a few minutes for each participant.

The training was also used to assure that the participant had good eye tracking, does not blink excessively, and can indeed adjust the length of her/his voluntary blinks to that of her/his spontaneous blinks.

### Experimental Setup

During the scanning sessions, participants lay supine in the scanner bore, their dominant eye being monitored by a video eye tracker (EyeLink 1000, SR research, Ontario, Canada) operating at a 500 Hz sampling rate. Each of the participants’ hands was placed on a separate response box: the right box was used to respond to the main experimental task, and the left box was used to trigger a self-initiated darkening when given an auditory cue (see Experimental Procedure below). Stimuli display was controlled by a PC running a custom Matlab script using Psychtoolbox-3 (Brainard 1997; Pelli 1997; Kleiner et al. 2007). The display was projected onto a screen positioned at the head end of the MRI scanner bore and viewed through a tilted mirror. All other light sources aside from the projector were shut or blocked, including LEDs and displays on the MRI scanner itself.

While several previous studies on eye blinks displayed black screens in order to mimic the retinal impact of blinks, this might be a rather crude control, depending on the response rate and contrast-ratio of the video projector used: Too slow response-rate would cause the darkenings to fade-in and out more slowly than intended; too low contrast-ratio would cause the black screens showed during the darkenings to be far brighter than true black and hence they may still illuminate the white surface of the MRI bore. This latter issue might lead to a systematic difference in the input to the retinal periphery between blinks (in which the periphery is truly occluded) and external darkenings (in which the MRI bore stays visible).

Hence, to better mimic the retinal darkening induced by blinks, we used an electro-optical LCD shutter (LC-Tech FOS series, model X-FOS(G2), LC-Tec Displays AB, Borlänge, Sweden) with a reaction time of 2-3 ms and 1800:1 contrast ratio. The shutter was placed in front of the projector’s lens (1B). Activation of the shutter was controlled by digital pulses sent from the experiment PC’s parallel port and converted into an analog driving voltage via a custom-built controller. When in closed (active) mode, the shutter turns from semi-transparent to opaque black. The extent of light blocking depends on the input voltage. Since the eyelid does not extinguish light transmittance entirely, we aimed to match the light transmittance of the shutter in its closed state to that of the eyelid. In a previous study, Ando and Kripke (1996) evaluated eyelid transmission with a visual threshold response. The estimated light transmission through the eyelids was 0.3% for blue and green, and 5.6% for red. Since the shutter could not differentially control transmittance for different wavelengths, we presented our stimuli in a green-scale only and set the shutter driving voltage such that it transmits 0.3% of the light in its maximally closed state.

**Figure 1.**
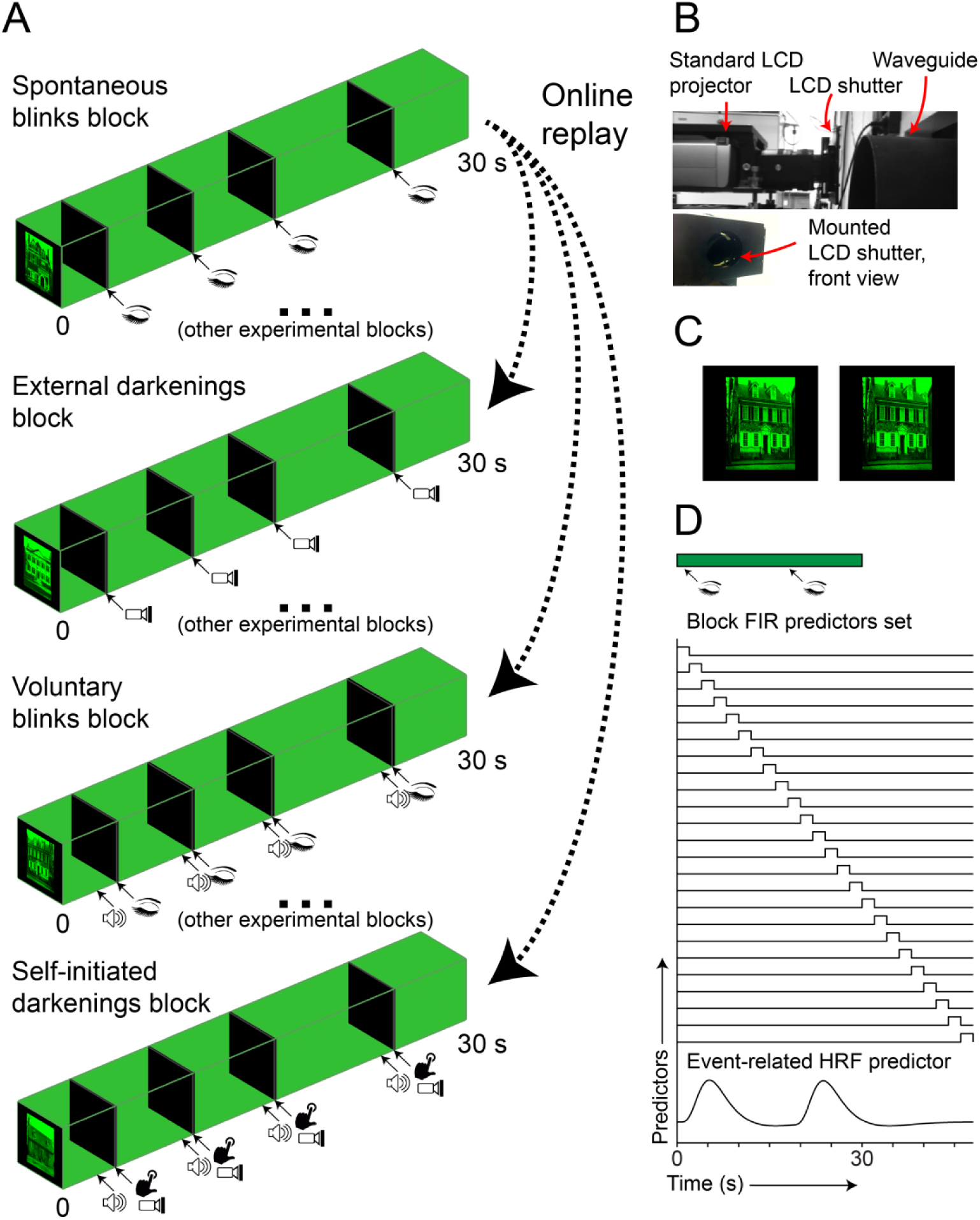
Experimental design. **(a)** Four experimental blocks **–** a spontaneous blinks block (top) and three blocks replaying its recorded blink latencies: external darkenings blocks, voluntary blinks block and a self-induced darkenings block. This particular block quartet was extracted from one of the participants’ experimental protocol. In the external darkenings blocks, the electro-optical shutter was shut according to the timings of replayed blinks. In voluntary-blinks blocks, an auditory cue preceding the replayed blinks’ latencies indicated the participant to blink. In the self-induced darkenings blocks, another auditory cue indicated the participant to press a button, which immediately triggered a darkening. Note that the replayed blocks are not consecutive and may occur in different runs, many minutes apart. **(b)** Electro-optical shutter installation. The shutter was placed between the lens of the video projector and the waveguide. **(c)** Visual discrimination task - an example house image and a non-identical probe image (note the chimney on the upper-right). **(d)** Mixed block-event GLM design matrix: example model of one block with two occurrences of a blink. An FIR predictor set (a pulse per each TR) accounts for the block-related response whereas an HRF-convolved event-related predictor accounts for the blinks’ (or other visual interruption) contribution to the BOLD signal. The green bar at the top marks the stimulus presentation time (0-30 seconds).

While we could not reproduce the gradual vertical covering and uncovering of the pupil during eye blinks, we simulated the gradual decrease and increase in incoming illumination by driving the voltage gradually from fully open to fully closed state and back with 10 ms long linear ramps.

### MRI acquisition

Participants were scanned with a Siemens 3T TrioMRI scanner equipped with a birdcage head coil used for RF transmit and receive. Blood oxygenation level dependent (BOLD) contrast was obtained using a T2* sensitive echo planar imaging (EPI): TR 2000ms; TE 30ms; flip angle 75°; field of view 256mm; voxel size 3x3x3 mm; 32 slices). For each participant, one volume of a T1 weighted high resolution (1x1x1 mm) anatomical image was acquired with a T1 weighted 3D-MPRAGE pulse sequence.

### Experimental procedure

A scheme of the experimental procedure is shown in Figure 1. Each run (8 runs in total per session) lasted 12 to 15 minutes and consisted of 30 seconds long blocks during which participants were instructed to focus their gaze on the center of a still image marked by a fixation cross. On each block, a single image of either a house or a face was presented for the entire 30 seconds period. Images subtended 10 x 8 visual angles (vertical x horizontal) and were presented in the center of a black display. The face images had a mean intensity value (out of 255 intensity levels) of 128.0±30.4 and RMS intensity contrast of 55.9±13.4, and the house images had a mean intensity value of 133.5±14.0 and mean RMS contrast of 88.8±7.2. To assure participants would stay attentive to the images during the blocks, they were instructed to perform a visual discrimination task, which was unrelated to our research question: 70% of the blocks were followed by a probe image in which a similar or an identical image to that presented during the block was shown. This probe image appeared after two seconds from block termination and was displayed until a response was collected (no more than four seconds response time was allowed). For faces, a similar image was a photograph of an identical twin of the person whose face was just presented, taken with the same pose. For houses, the similar images were produced by editing the originals in Adobe Photoshop to introduce slight feature-level changes (e.g., removing a window, see Figure 1C). The participant had to respond whether the probe image was same or different from the one presented during the block by pressing on one of two buttons on the right-hand response box. The participants’ responses were followed by auditory feedback, and a monetary reward was given for correct answers. The offset of the probe image (or the offset of the block, in trials in which no probe image was presented) and the next block’s onset were separated by 6, 6.5, 7, 7.5 or 8 seconds, randomly selected per block.

Across blocks, two factors were manipulated: stimulus category (whether it was a face or a house image) and visual interruption type (spontaneous blinks, cued voluntary blinks, unpredictable external darkenings and cued self-initiated darkenings). Only one type of visual interruption was introduced during a given block, resulting in a total of eight block conditions (2 stimulus categories x 4 visual interruption types). Obviously, spontaneous blinks kept also occurring during non-spontaneous-blinks blocks. This was addressed by the GLM analysis (see below). The number of blocks in each run was set to be 16 (four blocks for each visual interruption type, divided equally between houses and faces and presented in random order).

During spontaneous-blinks blocks, participants watched the still images with no visual interruption introduced other than their own spontaneous blinks, which were recorded and analyzed online using a semi-automatic approach (see ‘blink detection and duration measurement’ below). For each such a block, we recorded one ‘spontaneous blinks pattern’ (a set of blinks’ onsets and offsets relative to the block’s onset, excluding any blinks in the first 850 ms of the block). This pattern was then replayed three times in blocks of the same stimulus category: once as a cued voluntary blinks block, once as an (unpredictable) external darkenings block and once as a cued self-initiated external darkenings block. During cued voluntary blinks block, an auditory cue to blink was sounded 800 ms prior to the latency of each spontaneous blink in the replayed blink pattern. Similarly, during cued self-initiated darkenings blocks, an auditory cue to press a left-hand button was sounded 800 ms prior to the replayed blink latency. The button press immediately resulted in activating the shutter for a duration matched to that of the replayed blink. Last, during (unpredictable) external darkening blocks, the shutter was activated at the original replayed blinks’ latencies and durations. The recorded blink patterns were not replayed immediately but pushed into a queue instead. In each block, the blink pattern that was least recently recorded or replayed was pulled from the queue. The queue was maintained across runs (but not across sessions). While it is theoretically possible that the participants memorized the blink patterns, the replays of the same pattern were separated by several minutes, and hence we view such implicit or explicit memorization as unlikely (and indeed it was inconsistent with the participants’ reaction times, see Results). Auditory cues for voluntary blinking and self-initiated darkenings were counter-balanced between participants (a flute note and a hi-hat cymbals sound were used).

### Blink detection and blink duration measurement

While the Eye Link 1000 eye tracker automatically detects eye blinks, it sometimes mistakes short tracking losses to be eye blinks. Therefore, we used a semi-automatic approach for blink detection. During the fMRI scans, the experimenter monitored each spontaneous blinks block’s 30 s long pupil-size trace on a second linked workstation. The experimenter approved or rejected the detected blinks according to the presence of a gradual decrease and increase in pupil size (blink-unrelated tracking loss shows abrupt signal changes) and in some cases marked blinks that failed to be automatically detected by the eye tracker. In cases where the experimenter rejected an entire block due to lack of spontaneous blinks or due to eye tracking issues, another spontaneous-blinks-block was automatically introduced. In some runs, this caused the scanning sequence maximal duration (15 minutes) to elapse before 16 valid blocks were collected. However, the post hoc GLM modeling (see below) kept the analyzed blocks matched (in terms of blink patterns) and balanced (in terms of the four block conditions). Blinks onsets and offsets were determined according to the moment of maximal acceleration during pupil size reduction and maximal deceleration during pupil size increase, respectively. These two pertain to moments in which the eyelid touches the upper edge of the pupil, on its way down and on its way back.

Once an experimental session was over, the entire pupil-size timecourse for all runs was semi-automatically inspected for detecting blinks, and the results of this analysis were used for the offline analysis.

### fMRI data preprocessing

fMRI data were preprocessed using FEAT (fMRI Expert Analysis Tool) version 6.00, part of FSL (FMRIB’s Software Library, www.fmrib.ox.ac.uk/fsl). The following pre-statistics processing was applied: motion correction using MCFLIRT (Jenkinson et al. 2002) with each participant’s first run’s 100^th^ volume as a reference; slice-timing correction using Fourier-space time-series phase-shifting; non-brain removal using BET (Smith 2002); grand-mean intensity normalization of the entire 4D dataset by a single multiplicative factor; high-pass temporal filtering (Gaussian-weighted least-squares straight line fitting, with sigma=60.0s). The reference functional image of each participant was then co-registered to the participant’s anatomical volume using FLIRT (Jenkinson and Smith 2001; Jenkinson *et al.* 2002).

The anatomical image was processed using FreeSurfer 5.3 (<u>surfer.nmr.mgh.harvard.edu</u>) to obtain volumetric segmentation and cortical reconstruction (Dale et al. 1999; Fischl et al. 1999), which are necessary for cortex-based group analysis.

### First-level General Linear Model

For each participant, a general linear model was fit to his or her entire sequence of functional volumes. We used a mixed block-event design modeling approach, disentangling the contribution of the 30 s long face or house image presentation from the contribution of the visual interruptions (blinks and external darkenings). Since previous works have shown that the hemodynamic response to an image displayed for a prolonged interval exhibits a gradual decrease in signal over time (Grill-Spector et al. 1999; Gilaie-Dotan et al. 2008), modeling the visual response to images by a boxcar convolved with a hemodynamic response function (HRF) is likely to misspecify the hemodynamic response. Hence, we modeled blocks by a Finite Impulse Response (FIR) basis, accounting for the block-related response in each TR independently. The contribution of the visual interruptions on top of the block-related response was modeled by event-related HRF-convolved predictors (See 1D).

Face and house blocks were modeled separately, each by an FIR basis of 25 pulses, spanning from 0 to 48 seconds post block onset. Spontaneous blinks, voluntary blinks, self-initiated darkenings, and (unpredictable) external darkenings were each modeled by two event-related predictors, one for events occurring during face blocks and another for events occurring during house blocks (hence a total of eight event-related predictors of interest). Due to their collinearity with voluntary blinks and self-initiated darkenings, the auditory cues were not modeled by separate event-related predictors. Each event-related predictor was formed by placing a one-TR long pulse at each event’s onset and convolving the resulting timecourse with an HRF. We used the double gamma HRF model with the parameters estimated by Glover (1999). Event-related predictors of no-interest accounted for spontaneous blinks occurring during non-spontaneous-blinks-blocks, spontaneous blinks occurring between the blocks, probe images and button press responses to these probe images. A constant predictor accounted for baseline activity level.

Importantly, we only included in the GLM analysis blocks of fully replayed blink patterns, i.e., only quartets of blocks of each of the four conditions (spontaneous blink, voluntary blinks, external darkenings, and self-initiated darkenings) that used the same event latencies set. Since this matching and balancing was done across runs, we did not use the common FSL strategy of fitting a separate GLM model to each run. Instead, the GLM was fit to a concatenated file containing each participant’s entire sequence of functional volumes. This was done with a Matlab implementation of ordinary-least-squares which could accommodate the large memory required for fitting a GLM model to the entire concatenated file. Potential BOLD signal scale mismatches between runs were removed prior to concatenation by dividing each voxel by its within-run temporal mean. In the concatenated timecourse, each voxel’s grand-average BOLD signal level was restored by multiplication.

### Second-level statistical testing

Individual whole-brain maps of parameter estimates (PEs, ‘betas’) were projected by FreeSurfer to a template cortical mesh (‘fsaverage’) through cortex based alignment and then spatially smoothed using a 6 mm FWHM surface-based Gaussian filter (Hagler et al. 2006). For each contrast of parameter estimates (COPE), we obtained a whole-brain surface-map of its mean percent signal change value, as well as a t-value map, by averaging across the 13 participants.

Since Random Field Theory based cluster testing of fMRI effects has been recently shown to be optimistically biased (Eklund et al. 2016), our statistical testing was based on a non-parametric approach. For each grand-average COPE, an empirical null distribution was obtained by flipping the signs of individual-subject COPEs and re-computing the corresponding t-value whole-brain map. Since we tested 13 participants, simulating the entire null distribution (2^13^, 8192 different sign flipping combinations) was computationally feasible, and no randomized resampling was needed. Note that for binary contrasts, this sign flipping procedure is equivalent to permuting the conditions’ labels. For each simulated null t-map, we applied an in-house surface-based implementation of Threshold-Free Cluster Enhancement (TFCE, Smith and Nichols 2009), using the default parameters of E=0.5 and H=2. The TFCE transformation enhances vertices residing within clusters without having to explicitly define clusters by applying an arbitrary cluster-defining threshold.

The null TFCE maps were compared to the empirical TFCE map derived from the original COPEs (i.e., without sign-flipping) to assign each vertex with a p-value. We used a resampling-based false-discovery rate (FDR) control borrowed from Bioinformatics (Storey and Tibshirani 2003; Millstein and Volfson 2013):

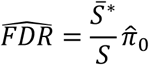

For any arbitrary TFCE threshold, 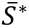 is the mean number of vertices that passed that threshold in each simulation of H0, and S is the number of vertices that passed the threshold in the true, empirical map. 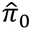 is the estimated proportion of true null hypotheses. Since the method of correctly estimating 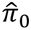 is still debated, we replaced 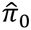 with 1, forming a slightly more conservative FDR estimate. Finally, we picked the lowest TFCE threshold that still held 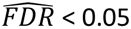 as our corrected statistical threshold. Unlike the Benjamini-Hochberg procedure (Benjamini and Hochberg 1995), this method does not require first computing uncorrected p-values but derives an FDR-controlling threshold directly from the data.

### Regions of interest definition

To define retinotopic regions of interest, we used a surface-based probabilistic atlas based on retinotopic mappings in multiple participants (Wang et al. 2014).

For category-preferring regions of interest, we used an individual, functional ROI definition. For each experimental block, we extracted the mean BOLD signal between four and eight seconds after the block’s onset. Then, within each participant, we contrasted face stimuli blocks and house stimuli blocks (regardless of the blocks’ visual interruption condition) by an unpaired two-sample t-test. An individual Fusiform Face-area (FFA, Kanwisher et al. 1997) ROI was defined by intersecting this t-map at t > 2.5 with a macro-anatomical Fusiform Gyrus surface-based label (Desikan et al. 2006), within each hemisphere. We then picked the largest contiguous cluster of above-threshold vertices. Similarly, a Parahippocampal Place Area (PPA, Epstein and Kanwisher 1998) was defined by intersecting the other tail of this t-map at t < -2.5 with a probabilistic functional PPA label (Weiner et al. 2017) and again, selecting the largest contiguous cluster.

For all ROIs, the responses were sampled in left and right hemispheres and were averaged together with equal weight.

For group-level whole-brain maps visualization purposes, we generated an FFA ROI by a second-level contrast between all faces blocks with all house blocks (using a 30 s HRF-convolved boxcar block-model). We thresholded the resulting t-map with a threshold of t > 3, intersected the result with the Fusiform Gyrus surface-based label and selected the largest contiguous patch in each hemisphere. Group-level PPA coincided with PPA-1 and PPA-2 of Wang and colleagues’ atlas participants (Wang *et al.* 2014) and hence was not drawn.

### Blink duration matching control

To further control for potential duration differences between spontaneous and voluntary blinks, we matched the durations of the two blink types selecting a subset of spontaneous blink events and a subset of voluntary blink events so the two subsets will have a highly compatible duration distribution. This was done using an iterative procedure (Golan *et al.* 2016): in each iteration, a pair of a spontaneous blink and a voluntary blink was added to the selected subsets. The criteria for selection were a duration difference of no more than 5 ms between the spontaneous and voluntary blink, and the minimization of the mean duration of the two subsets. This was repeated until no eligible pairs were left for selection.

The matching procedure reduced the number of modeled blinks (spontaneous and voluntary) to 34.3-74.7% (M=58.5%) of the original number of blink events per subject, resulting in 29.0-520.0 (M=286.8) duration-matched blink-pairs per subject. For this procedure, experimental blocks that were not fully replayed were not excluded. Unselected subsets of spontaneous and voluntary blinks were modeled by predictors of no interest, hence effectively removed from the GLM analysis.

## Results

All 13 participants who underwent the fMRI experiment completed short preliminary training in generating voluntary blinks of a duration similar (±1 SD) to their spontaneous blinks’ mean duration. During the fMRI experiment, the participants viewed photographs of either faces or houses, with each image presented continuously throughout a single 30-second long experimental block. The performance in the discrimination task, comparing a probe image appearing after 70% of the experimental blocks to the image displayed during the preceding block, was 91.0%±6.3% for faces and 69.4%±12.7% for houses.

In about one-quarter of the experimental blocks, no additional manipulation was introduced, and the participants’ spontaneous blinks were monitored online. For each such ‘spontaneous blinks block’, the set of the blinks’ onsets and offsets relative to the block’s onset was used to generate three yoked matching experimental blocks which were presented later in the experimental session (see 1A and Materials and Methods). In ‘external darkenings’ blocks, an LCD shutter (1B) was used to simulate the retinal impact of spontaneous blinks by whole-field darkenings of matched latencies and durations. In ‘self-initiated darkenings’ blocks, the same darkenings were triggered by an auditorily cued button press. Last, in the voluntary blinks blocks, the participants responded to another auditory cue with a voluntary blink. Both auditory cues were presented 800 ms before the mirrored onsets of the yoked spontaneous blinks block, to compensate for the participants’ expected reaction time. Compliance to both cues was high (M=90.4%, SE=1.1% for self-initiated darkenings and M=90.9%, SE=1.6% for voluntary blinks). Supplementary Figure 1 presents the latency (in relation to block onset) distributions of the four types of visual interruptions. Supplementary Figure 2 presents their duration distributions. Note that due to the high compliance with the task, the latency distributions of the four conditions within a given participant were very similar.

**Figure 2.**
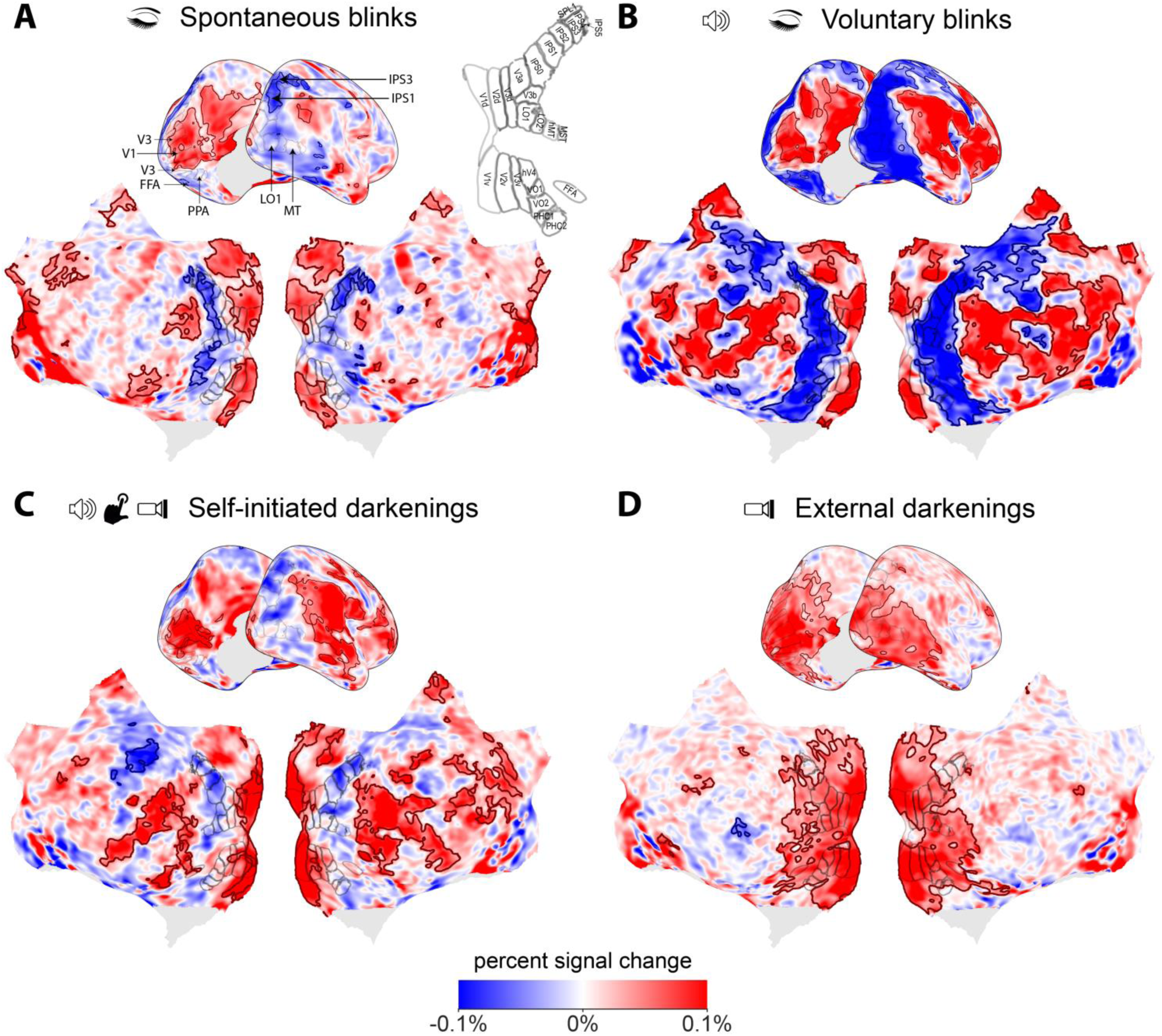
Hemodynamic response whole-cortex maps to (A) spontaneous blinks, (B) voluntary blinks, (C) self-initiated darkenings, and (D) (unpredictable) external darkenings. The color scale denotes BOLD percent signal change (normalized within each vertex by the constant predictor parameter estimate). Red or blue outlines, respectively mark significant activity-increases or -decreases related to the four events (*p*_FDR_ < 0.05 for all vertices within the outlines *-* tested non-parametrically on threshold-free cluster-enhanced images). The thin black outlines trace the borders between retinotopic regions from Wang and colleagues’ (2014) surface-based functional atlas, as well as the Fusiform Face Area (FFA). Note the activation of the early visual cortex by all four conditions and in contrast, the differential effect in higher-level visual cortex: deactivation by spontaneous blinks, voluntary blinks, and self-initiated darkenings, and a positive activation in response to unpredictable external darkenings.

To confirm that no memorization of replayed blink patterns took place, we computed the mean reaction time to voluntary blink cues in two different conditions: when the particular blink pattern was replayed for the first time, and when the particular blink pattern was already replayed twice, in a preceding self-initiated darkenings block and in a preceding external darkening block. Inconsistent with memorization of blink patterns, the participants were in fact nominally (but non-significantly) slower in the third exposure (M=879.9 ms, SE=46.3 ms) than in the first exposure (M=869.5, SE=43.6 ms; paired-samples t-test, t(12)=0.68, *p* = 0.51). For cues to self-initiate a darkening, the participants were slightly faster in the third exposure (M=1047.6 ms, SE=36.4 ms) than in the first exposure (M=1076.0 ms, SE= SE=44.3 ms) but this difference was not significant as well (paired-samples t-test, t(11)=1.22, *p* = 0.25, the reduced degrees of freedom is due to one participant for which the manual RT was not logged).

Using a mixed event/block GLM (Figure 1D), we estimated the contribution of the event-related visual interruptions (spontaneous blinks, voluntary blinks, external darkenings, and self-initiated darkenings) to the BOLD signal, after accounting for the hemodynamic response to the still images presented throughout the blocks. Figure 2 presents whole-cortex maps of the event-related response to the four visual interruptions normalized to percent signal change (normalized within each vertex by the constant-predictor parameter-estimate, which reflects the baseline signal of the vertex after accounting for all other predictors). To avoid the imager’s fallacy (Henson 2005), we show the entire data, and instead of thresholding the map, significant effects are indicated with red and blue outlines.

Spontaneous blinks (Figure 2A) showed a significant event-related BOLD increase in the representation of the retinal periphery of V1-V3 and a small but widespread deactivation of mid- and high- level visual regions, (there were further significant activations and deactivations in non-visual regions, which are beyond the scope of this report). Voluntary blinks showed a similar activity pattern, with a more pronounced and extensive deactivation of high-order visual regions (Figure 2B). Formally contrasting the two blink types (3B) revealed that the activity reduction related to voluntary blinks was indeed significantly greater than that related to spontaneous blinks (*p*_FDR_ < 0.05, here and in the following contrasts reported as significant), an effect spanning both ventral and dorsal high-level visual regions. It should be reiterated that these response estimates were acquired while accounting for the far larger responses to the visual stimuli by a separate set of FIR predictors. Hence, this mild (~0.1% signal change) negative response to blinks in high-level visual cortex should not be interpreted as an activity decrease below blank screen level; Rather, it signifies an activity decrease below the activity levels expected had the face/house stimulus been viewed without any visual interruption.

**Figure 3.**
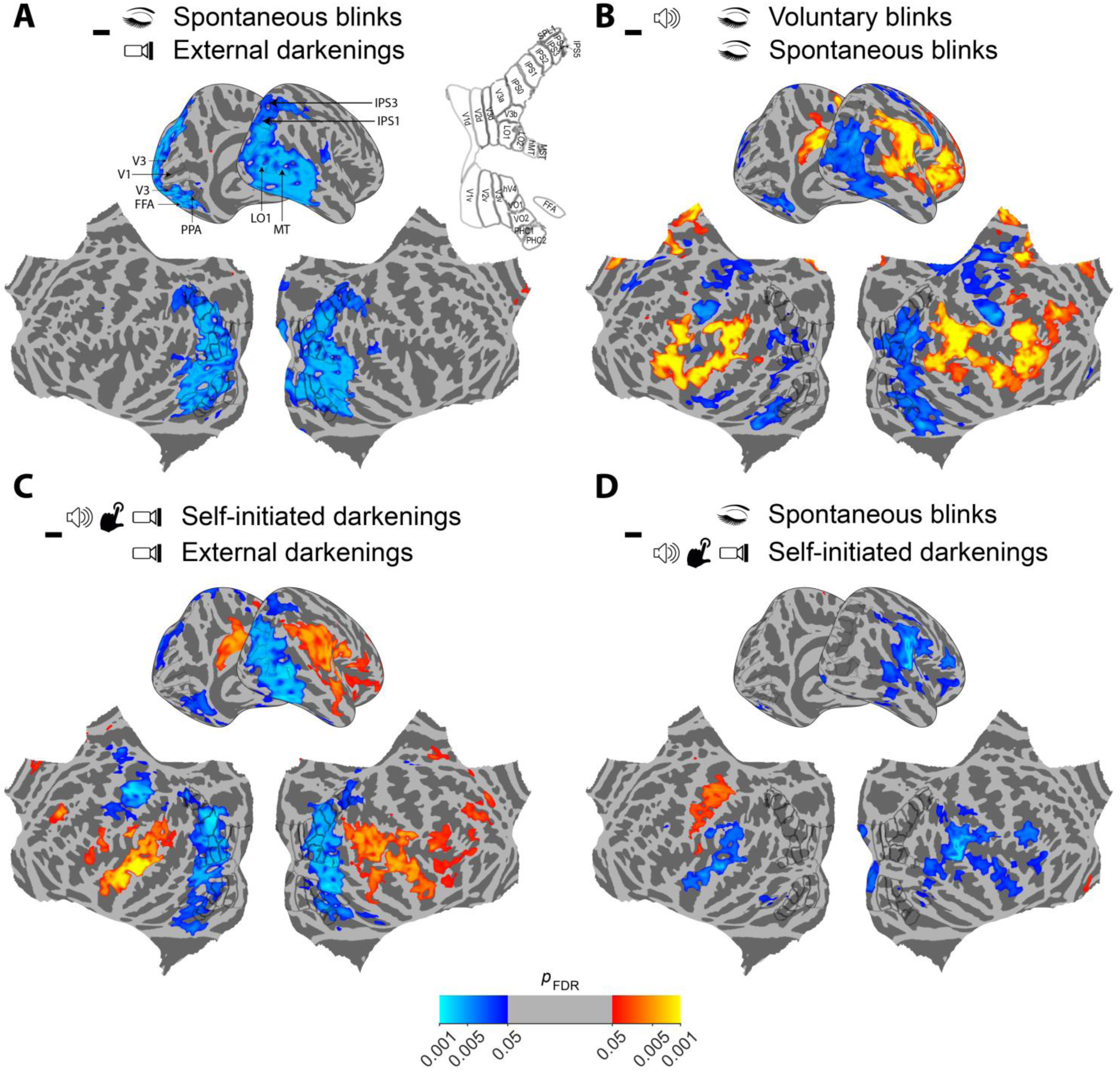
Pairwise contrasts of the different visual interruptions. The color scale is logarithmically mapped to the vertices’ FDR-corrected p-values (tested on TFCE maps). Red-yellow hues mark positive effects, and blue-cyan hues mark negative effects. Notable effects: (A) Significantly more negative response in mid- and high-level visual cortex to spontaneous blinks compared with external darkenings. (B) Significantly more negative response in mid- and high-level visual cortex to voluntary blinks compared with spontaneous blinks. (C) Significantly more negative response in mid- and high-level visual cortex to self-initiated darkenings compared with spontaneous blinks. (D) Similar (i.e., not statistically distinguishable) mid- and high-level visual cortex responses to spontaneous blinks and self-initiated darkenings.

The pattern of visual responses to self-initiated darkenings (Figure 2C) was similar to that of spontaneous blinks, yet the activation reduction in higher-level visual regions did not cross the significance threshold for this condition (but see the ROI analysis, Figure 4, for un-thresholded responses). Contrasting self-initiated darkenings with spontaneous blinks found that self-initiated darkenings were not statistically distinguishable from spontaneous blinks in the high-level visual cortex (Figure 3D). The two conditions significantly differed mostly in auditory and motor cortical regions, reflecting the impact of the auditory cue and manual motor response in the self-initiated darkenings condition.

**Figure 4.**
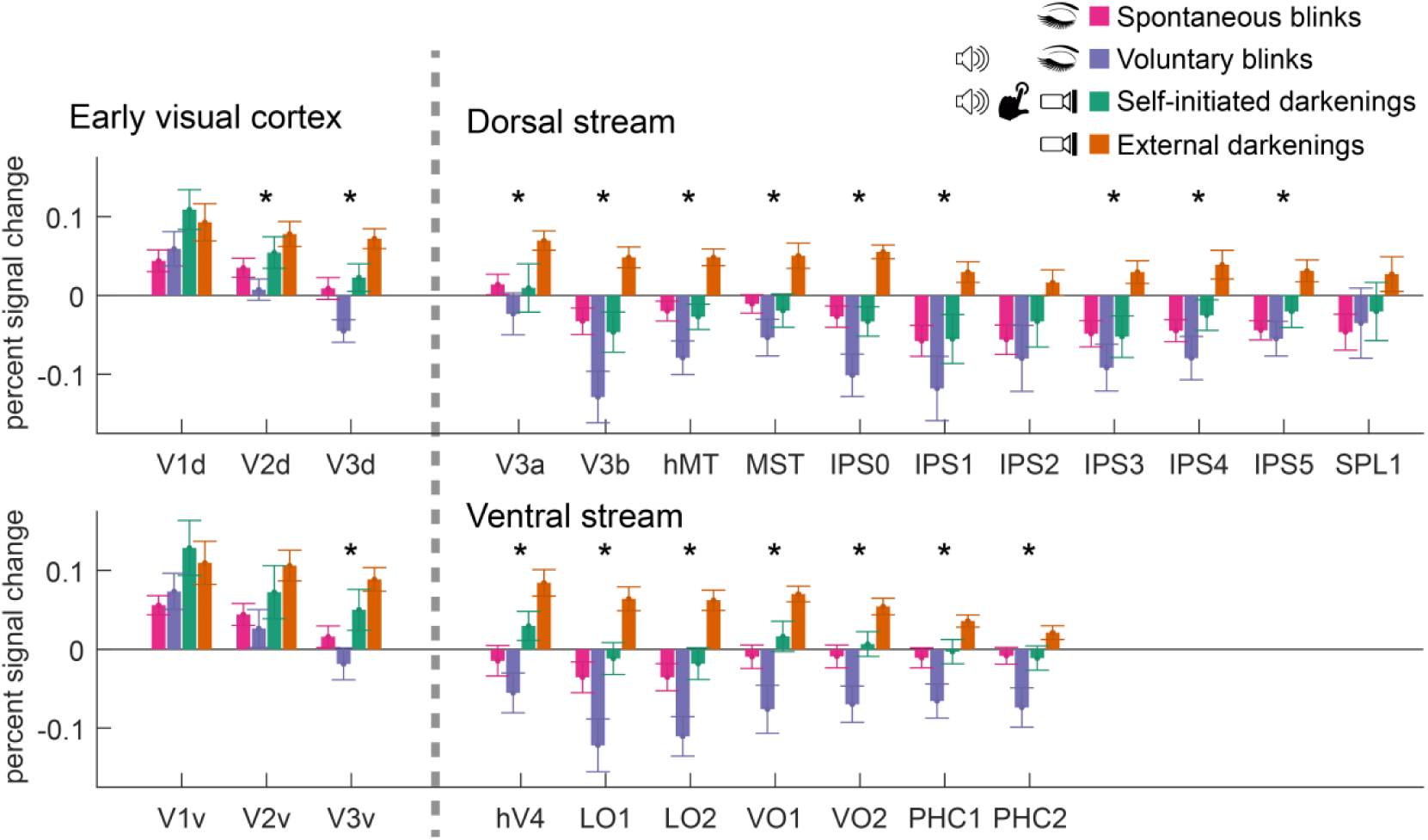
Parameter estimates of the responses to spontaneous blinks, voluntary blinks, self-initiated darkenings, and external darkenings. Asterisks denote a significant (*p*_FDR_ < 0.05) main effect of visual interruption type within a particular region of interest. Error bars are ±1 SEM, reflecting between-subjects variability. For analysis within the FFA and PPA regions, see Supplementary Figure 4).

Last, external darkenings (Figure 2D) showed a different response pattern, with mild but widespread activity increase in both low, mid- and high-level regions, with no trace of activity reduction in visual areas. In mid- and high-level visual cortex, this response was significantly more positive than the responses related to spontaneous blinks (Figure 3A), voluntary blinks (not displayed) or self-initiated darkenings (Figure 3C).

In order to compare the hemodynamic response patterns in different visual regions to the four conditions across the visual hierarchy, we sampled the estimated responses using 24 retinotopic regions defined by a probabilistic functional atlas (Wang *et al.* 2014). The resulting ROI-sampled data (Figure 4) showed an increasing suppression of the responses to spontaneous blinks, voluntary blinks, and self-initiated darkenings, an effect gradually appearing and intensifying across the visual hierarchy. A repeated measures ANOVA with the factors of ROI and visual interruption (spontaneous blink/voluntary blink/self-initiated darkening/external darkening) found a significant ROI x visual interruption type interaction (*F*(69,828) = 2.76, *p* = 0.0184, Greenhouse-Geisser corrected), as well as significant main effects of ROI and visual interruption type (both *p* < 0.001, Greenhouse-Geisser corrected). Testing for a visual-interruption-type effect within each ROI found significant effects in dorsal V2 & V3, V3a, hMT, MST, IPS0, IPS1, IPS3, IPS4, IPS5, as well as ventral stream regions – ventral V3, LO1, LO2, VO1, VO2, PHC1, PHC2 (*p*_FDR_ < 0.05, Greenhouse-Geisser corrected). We did not test the simple effects within ROIs as these were already assessed in the whole-cortex contrasts shown in Figure 3.

### Controlling for potential duration mismatches

Can the observed differences between the visual-interruption conditions be explained by differences in retinal blackout duration? While the self-initiated and external darkenings’ durations were matched to the spontaneous blinks’ durations by our online replay procedure, the duration of the voluntary blinks depended on the participants’ performance. The participants underwent short training aimed at minimizing this issue, but some of them still produced spontaneous and voluntary blinks of non-identical duration distributions during the main experiment (Supplementary Figure 2). Could these mismatched events explain the greater reduction in high-level visual cortex activity in response to voluntary blinks compared with spontaneous blinks? To test that, we ran the GLM model estimation after selecting a subset of blink events matching the duration distributions of spontaneous and voluntary blinks (Supplementary Figure 3A). The responses to the duration-matched spontaneous blinks (Supplementary Figure 3B) and duration-matched voluntary blinks (Supplementary Figure 3C) still clearly showed a significantly greater deactivation for voluntary blinks than spontaneous blinks (Supplementary Figure 3D). Hence, duration differences could not explain the greater deactivation of high-level visual cortex related to voluntary blinks compared with spontaneous blinks.

### Testing for interactions with stimulus category

In Golan and colleagues (2016), intracranial-EEG responses in category-selective regions showed an interaction of stimulus category with visual interruption type. Specifically, electrodes in face-selective regions (such as the Fusiform Face Area, FFA) showed significant activity increases in response to the reappearance of face images but not to the reappearance of non-face images, and this selective reappearance-related response occurred when the reappearance was visible, following an externally induced blank, but not when it was invisible, following a spontaneous eye blink. Thus, we designed the current experiment to allow the assessment of this effect in fMRI by the inclusion of face and house blocks.

However, contrasting each type of visual interruption (spontaneous blinks, voluntary blinks, self-initiated darkenings or external darkenings) between the face and house blocks failed to reveal any region passing the whole-cortex maps’ statistical threshold. Contrasting the external darkenings minus spontaneous blink effect between the face and house blocks did not result in any significant category-selective activation as well. To ease the multiple comparisons problem and any remaining inter-subject anatomical variability, we turned to an ROI-based approach, defining individual-level functional FFA and PPA regions of interest (See Materials and Methods). Within each ROI, we measured the responses to the four visual interruption conditions for each stimulus type (face/house, Supplementary Figure 4). There was a significant effect of condition (F(3,36) = 7.54, p = 0.005, Greenhouse-Geisser corrected), but the other ANOVA effects, including the triple interaction of stimulus type x visual interruption type x ROI, were not significant. Furthermore, while the responses to external darkenings in the PPA were indeed greater during house blocks, this was true also for external darkenings occurring during face blocks. Hence, we failed to observe the category-selective positive response to (the termination of) external stimulus omission, which we previously observed in intracranial EEG (2016). This seeming discrepancy between intracranial EEG and fMRI signals might be related to temporal resolution limitations of the BOLD signal (see Discussion below).

## Discussion

Using a mixed block/event design fMRI experiment, we found that: (a) spontaneous blinks, voluntary blinks and external darkenings (either surprising or self-initiated) activated the peripheral fields of V1 to V3. The blink-related early visual activation has been observed in fMRI before (e.g., Hupé *et al.* 2012), however the finding that the same response is also triggered by external darkenings strongly argues for a retinal origin of this effect. (b) In mid- and high-level visual cortex, a considerably different pattern of response was observed: while unpredictable external darkenings positively activated the entire visual hierarchy, predictable events of different kinds (i.e., spontaneous blinks, voluntary blinks, and self-initiated darkenings) induced a slight (but significant and widespread) reduction of the BOLD visual response. This effect was more pronounced in the dorsal stream (see Figure 4, top panel). The blink-related activity reduction is consistent with and generalizes previous findings in voluntary blinks over checkerboard stimuli (Bristow, Frith, *et al.* 2005). However, the finding that self-initiated darkenings cause a similar activity reduction offers a new interpretation of the suppressive response to blinks, suggesting the involvement of a general-purpose prediction-related mechanism in addition to the specialized ‘hardwired’ oculomotor efferent copy. Last, voluntary blinks showed the strongest negative modulation of higher-level visual cortex activity, even when their durations were equated to those of spontaneous blinks by post hoc selection.

### Interpreting the BOLD responses in light of previous electrophysiological findings

Interpreting the current results requires caution due to the sluggish nature of the BOLD signal compared with the relatively brief duration of blinks. If we had the fMRI results alone, the most straightforward interpretation of the mid- and high-level visual cortex effects would have been that unpredictable external darkenings excite the high-level visual cortex whereas spontaneous and voluntary blinks, as well as self-initiated external darkenings, reduce its activity. However, time-resolved iEEG recordings acquired across the visual cortex during a similar task (Golan *et al.* 2016) mandate a more nuanced interpretation: these recordings showed that the unpredictable darkenings caused a small, transient activity drop (see Golan et al., 2016, Figure 5—figure supplement 1b), which was followed by a far larger positive overshoot associated with picture reappearance. Here, the BOLD signal summates these two components, leading to a net positive hemodynamic response in both low- and high- level visual cortex to unpredictable external darkenings. In contrast, for spontaneous and voluntary blinks, the positive overshoot that follows the activity reduction was shown in the intracranial data to be fully or partially suppressed in higher-level regions, likely leading to the net negative BOLD response observed here. Hence, we speculate that the net negative BOLD response to the self-initiated darkenings introduced in high-level visual cortex may reflect the combination of a bottom-up reduction in activity and a top-down suppression of the transient positive responses to the reappearing image. This top-down suppression was absent when the darkenings were triggered by the experimenter instead of the participants themselves.

### Non-specific prediction mechanisms and eye blinks

Suppression of blink-related visual responses is commonly viewed as a product of a specialized ‘hardwired’ efferent copy pathway, linking oculomotor activity with suppression of visual responses (e.g., in the seminal work by Volkmann *et al.* 1980; Volkmann et al. 1982). This view is also supported by psychophysical (Bidder and Tomlinson 1997) and electrophysiological (Golan et al. 2017) evidence linking blink suppression with saccadic suppression.

The current finding that self-initiated darkenings triggered a net negative high-level visual response, similarly to spontaneous eye blinks, suggests that such a hardwired pathway might not be the only extra-retinal mechanism enabling blink-related neural suppression. A prominent candidate for such a non-specific mechanism is the ability to flexibly predict sensory events. Under the predictive coding model, a cardinal part of the measured response reflects the difference between the predicted input and the actual one (i.e., prediction-error). Within this framework, the reduced activation in high-level visual cortex in all but the unpredictable darkening may be the result of the ability to predict the visual interruption and therefore reduce the prediction error signal. While an efferent copy is ultimately a predictive signal, it is a highly specialized one, and relies either on pervasive exposure or on dedicated, innate pathways. In contrast, humans and animals can readily learn to predict visual events from quickly learned repetitions and regularities and this rapidly learned prediction becomes increasingly manifested in the cortical response as sensory signals flow downstream (e.g., see Summerfield et al. 2008 for the cortical effect of predictability in recurring arbitrary sequences of visual stimuli ; Dürschmid et al. 2016 for a comparable effect in the auditory domain). The significant fMRI contrast of unpredictable versus self-initiated external darkenings (Figure 3C) indicates that such short-term and flexible prediction may affect high-level visual cortex even when the prediction is inter-modal (here, motor-visual), adding to the existing evidence pointing at this direction (e.g., reduced high-level category-selective responses to auditorly cued faces and houses, den Ouden et al. 2010).

However, flexible, ad hoc prediction does not fully explain the current results. First, according to this account, predictability should modulate the responses of category-selective regions only for stimuli of their preferred category, an interaction we did not observe. Second, voluntary blinks, even when precisely matched to spontaneous blinks in terms of duration, caused a stronger activity reduction in high-level visual cortex. A potential explanation for this discrepancy might be that voluntary blinks are suppressed by a superposition of two forms of predictive signals: the suppression of spontaneous blinks is conditioned on automatic motor events that occur thousands of times a day, whereas the suppression of the self-initiated darkenings is conditioned on voluntary events and is flexibly acquired, during the experiment. Cued voluntary blinks might be affected by both kinds of predictive signals, hence inducing a greater deactivation in high-level visual cortex.

To summarize, while our results do not rule out efferent-copy effects causing blink-related suppression of visual processing, they indicate that general predictive learning may explain at least some facets of how blinks affect the visual cortex. In light of this finding, it may be worthwhile to revisit the already published psychophysical, electrophysiological and fMRI experiments on eye blinks and replicate them with added predictability controls. Such an endeavor may shed further light on how general or specific blink suppression is.

## Acknowledgments

We would like to thank Fanny Attar, Nahum Stern and Edna Haran-Furman for assistance with fMRI scanning and Michal Harel for assistance with cortical reconstructions.

